# A human brain network specialized for abstract formal reasoning

**DOI:** 10.1101/2025.10.21.683445

**Authors:** Hope Kean, Alex Fung, Chiebuka Ohams, Jason Chen, Josh Rule, Joshua Tenenbaum, Steven Piantadosi, Evelina Fedorenko

**Affiliations:** MIT; Grossman Center for Quantitative Biology and Human Behavior at the University of Chicago; UC Berkeley

## Abstract

Humans stand out in the animal kingdom for their ability to reason in highly abstract ways. Using a deep-data precision fMRI approach, we identify and richly characterize a network of frontal brain areas that support abstract formal reasoning. This ‘abstract reasoning’ network robustly dissociates from the domain-general Multiple Demand network—the current leading candidate substrate of fluid intelligence—as well as from three other networks supporting high-level cognition: the language network, the intuitive physical reasoning network, and the social reasoning network. Finally, the areas of this network respond robustly during both deductive and inductive reasoning, during classic matrix reasoning problems, and when solving multiplication and division problems. This network may therefore support the most abstract forms of reasoning, possibly constituting a human-specific adaptation.

## Introduction

Abstract logical reasoning is the hallmark of human intelligence. From the construction of mathematical proofs to the framing of legal arguments and scientific theories, logical reasoning has powered almost every sphere of human achievement. Although other species rival or surpass us in domains like perception, motor control, hierarchical social inference, and both long-term and working memory (Cronin et al., 1989; Cheney & Seyfarth, 1990; Clayton & Dickinson, 1998; Connor et al., 2000; Inoue & Matsuzawa, 2007; Tobalske et al., 2007), humans stand out in their facility manipulating abstract symbolic structures, such as those found in logic or mathematics, detached from any particular knowledge domain or sensory modality. Although this is arguably the defining signature of our species’ intelligence, many questions remain about the cognitive, neural, genetic, and computational basis of abstract logical thought (Donald, 1993; 2001; Gentner et al., 2001; Penn et al., 2008; Pinker, 2010; Duncan et al., 2012; Tomasello, 2014; 2019; Heyes, 2018; Griffiths, 2020; Cantlon & Piantadosi, 2024).

The search for neural circuits underlying human fluid intelligence has converged on a distributed set of frontal and parietal brain regions often called the Multiple Demand (MD) network (Duncan, 2010; Mitchell et al., 2016; Assem et al., 2020). Functional MRI shows that these areas are recruited during diverse goal-directed behaviors, often probed with demanding executive tasks, such as working memory, inhibitory control, and task-switching paradigms (Duncan & Owen, 2000; Fedorenko et al., 2013; Hugdahl et al., 2015; Shashidhara et al., 2019; Assem et al., 2024). All these tasks elicit responses across the MD network with the more difficult conditions eliciting a stronger response (Duncan & Owen, 2000; Fedorenko et al., 2013; Shashidhara et al., 2019). The MD network is also engaged by certain forms of reasoning, including different kinds of mathematical reasoning (Monti et al., 2012; Fedorenko et al., 2013; Amalric & Dehaene, 2016, 2019) and understanding pieces of computer programs (Ivanova et al., 2020; Liu et al., 2020). Activity in this network scales with individual differences in working memory capacity and fluid intelligence (Gray et al., 2003; Lee et al., 2006; Basten et al., 2015; Hilger et al., 2017; Assem, Blank et al., 2020), and damage to its components—but not other parts of the brain—is associated with broad executive deficits and loss of fluid intelligence (Woolgar et al., 2010; Gläscher et al., 2010; Woolgar et al., 2018). Together, these functional and causal lines of evidence make the MD network the leading candidate substrate for domain-general human intelligence.

However, relatively few brain-imaging studies have directly targeted formal logical reasoning (Goel et al., 1997; 1998; 2000; Goel & Dolan, 2001; Christoff et al., 2001; Acuna et al., 2002; Knauff et al., 2002; 2003; Noveck et al 2004; Canessa et al., 2005; Fangmeier et al., 2006; Reverberi et al., 2007; 2009; Wendelken & Bunge, 2010; Cho et al., 2010; Monti & Osherson, 2012; Prado et al., 2010; 2013; Chuikova et al., 2025; for reviews, see Wharton & Graffman, 1998; Houdé & Tzourio-Mazoyer, 2003; Goel, 2007; Prado et al., 2011; Oaksford, 2015; Heit, 2015; Wertheim & Ragni, 2020; Wang et al., 2020). The ones that did used diverse paradigms and analytic approaches and yielded heterogeneous and sometimes conflicting findings (Osherson et al., 1998; Parsons & Osherson, 2001; Monti et al., 2007; Monti et al., 2009). Furthermore, most prior studies have not directly compared brain responses to logic tasks and other demanding tasks (cf. Coetzee & Monti, 2018; Ruff et al., 2003; Kroger et al., 2002; 2008) and have often relied on precarious reverse inference (Poldrack, 2006) to interpret the activations. These limitations aside, many brain structures that have been implicated in these earlier studies resemble parts of the MD network, but some studies also report areas that appear to fall outside of the MD network’s boundaries—in some cases in the anterior extent of the left frontal lobe (Christoff et al., 2001; Hampshire et al., 2011; Chuikova et al., 2025; see <u>Discussion</u>). These anterior-most frontal regions have also been implicated in aspects of higher-order cognition based on human lesion work (Milner, 1963; 1964; Luria 1966; Goel & Grafman, 1995; Duncan et al., 1996; Goel et al., 1997; Reverberi et al., 2005a; 2005b; 2005c; Roca et al., 2010; 2011), and highlighted as showing pronounced divergences between humans and non-human primates in comparative anatomical studies (Semendeferi et al., 2001; Neubert et al., 2014; Donahue et al., 2018; Bryant et al., 2025). In tandem, this evidence suggests that brain areas outside the Multiple Demand network may contribute to fluid intelligence and reasoning. Dissociations between certain reasoning tasks and executive abilities in behavioral studies of individual differences (Hampshire et al., 2012; Stanovich et al., 2011) further corroborate the idea that multiple systems may support abstract thought.

To illuminate the neural basis of logical thought, we used a paradigm that isolates the complexity of deductive inferences (Coetzee & Monti, 2018), which reliably engaged a set of bilateral frontal areas, including around the frontal pole, and then systematically characterized these areas across 10 fMRI experiments (all conducted in a set of 24 participants, each completing between one and three 2-3 hour scanning sessions), including examining their relationship with known cognitive systems. To foreshadow our critical findings, the brain areas that support logical reasoning robustly dissociate from the Multiple Demand network, support both deductive and inductive abstract reasoning, as well as aspects of mathematical reasoning, and are distinct from the language network, the intuitive physical reasoning network, and the social reasoning network. This newly identified brain network may therefore support the most abstract forms of human reasoning.

## Results

To identify brain areas sensitive to deductive reasoning complexity, we adopted a paradigm that uses verbal syllogisms in a classic three-sentence (two premises, one conclusion) format and contrasts harder and easier deductions (**Figure 1A**; Coetzee & Monti, 2018). The harder deductions require a backward, contrapositive inference (reasoning by Modus Tollens), whereas the easier deductions require forward, direct conditional inference (reasoning by Modus Ponens) (see <u>Methods</u> for details; all stimuli and experimental scripts are available at OSF: osf.io/hqa23). This complexity difference is evidenced behaviorally (Wason, 1968; Evans & Newstead, 1977; Johnson-Laird, 1983; 1999; Girotto et al., 1997; Barrouillet et al., 2000; Coetzee & Monti, 2018), including in our data (**Figure 1; Supp. Fig. 2**).

**Figure 1.**
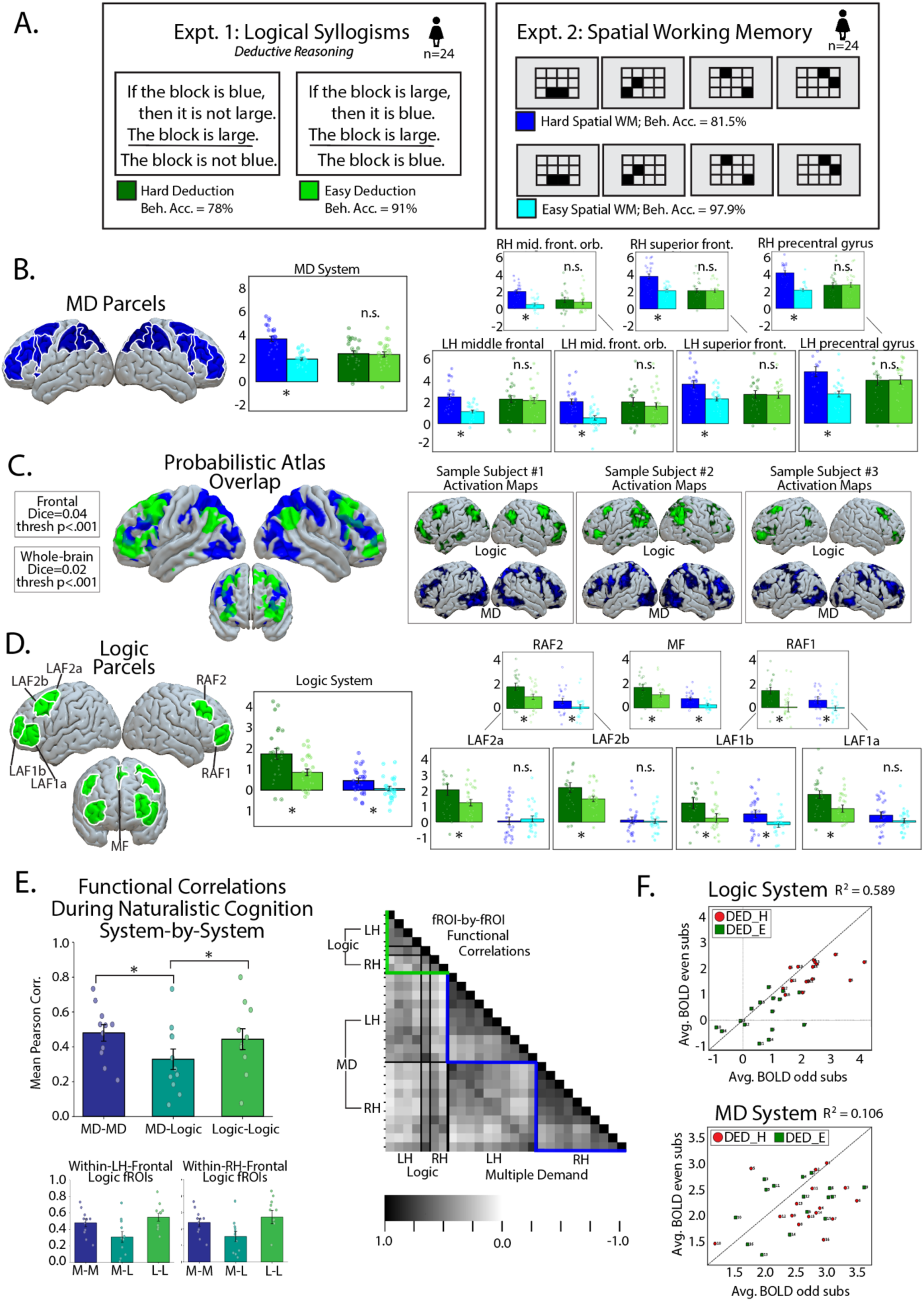
The logic system and its relationship with the Multiple Demand (MD) system. **A.** Experimental paradigms: participants performed a deductive reasoning task (Logical Syllogisms; Hard vs. Easy conditions) and a spatial working memory task (Hard vs. Easy). **B.** The MD system parcels and their responses to the Hard>Easy contrasts from both tasks, averaged across fROIs and broken down by individual fROIs. Logic reasoning elicited no reliable MD engagement, in contrast to the robust Hard>Easy effect for spatial WM. **C.** Probabilistic logic atlas (green) overlaid with the probabilistic MD atlas (blue) with a cut-off of 0.2. Boxes contain the dice overlap scores (p<0.001) between the individual activation landscapes for the logic task and spatial WM task in individual participants in both the left-frontal lobe and in the whole-brain. Sample individual activation maps demonstrate the distinct territories that the logic (green) and MD (blue) activations occupy in individual participants. **D.** Logic system parcels and the network-level as well as individual fROI responses to the logic localizer (Hard>Easy Deduction), averaged across fROIs and broken down by ROI. All regions showed reliable logic selectivity, with minimal response to spatial WM. **E.** Functional correlations during naturalistic cognition. logic areas were more strongly correlated with one another than with MD regions, and within-hemisphere logic ROIs showed reliable functional coupling. Right: pairwise correlation matrix across fROIs in the logic and MD systems. **F.** Split-half reliability of the BOLD response across even- and odd-indexed participants. The logic system showed high reliability across participants, whereas the MD system showed weaker and less consistent responses to the syllogisms.

To examine the relationship of logic-sensitive areas to the Multiple Demand (MD) network, we included a commonly used MD localizer—a spatial working memory task (Assem, Blank et al., 2020) (Experiment 2). To test whether the responses in the logic areas generalize to other forms of logical reasoning, we included three paradigms that all require abstract, structured inference: (i) syllogistic deductive reasoning on symbolic (non-verbal) problems, (ii) inductive reasoning (program induction: inferring rules from examples; Rule et al., 2020; 2024), and (iii) a task commonly used to measure IQ / fluid reasoning abilities, which requires a combination of deductive and inductive inferences (Cattell-type matrix problems; Woolgar et al., 2013) (Experiments 3-5). To examine the role of the logic areas in arithmetic—another, much more studied domain of formal reasoning (Dehaene, 1997)—we included three experiments spanning the four arithmetic operators (Experiments 6a-c). Finally, to examine the relationship of these areas with other known cognitive systems, we included localizers for the language network (Fedorenko et al., 2010, 2024), the intuitive physical reasoning network (Fischer et al., 2016), and the social reasoning network (Saxe & Kanwisher, 2003; Jacoby et al., 2016) (Experiments 7-10).

Twenty-four participants (mean age = 26.4 years, st. dev. = 8.6 years; 14 females; **Table 1**, <u>Methods</u>) were recruited for this deep-data study. Each participant performed the deductive reasoning task and between three and nine additional experiments (**Supp. Table 4** for details).

**Table 1.** The responses of the logic system to logic and non-logic contrasts at the network level, and the responses of the MD, Language, Intuitive Physics, and ToM systems to the logic contrast and their own localizer task contrast(s). The different systems correspond to different columns, and the different tasks (in Experiments 1-10, E1-E10; note that some tasks have multiple contrasts) correspond to rows. The statistics come from the linear mixed-effects models (see <u>Methods</u>; see Supp. Table 1 for the results broken down by fROI).

| Parameters |  | Logic Sys |  | MD Sys |  | Lang Sys |  | Int. Phys. Sys |  | ToM Sys |  |
| --- | --- | --- | --- | --- | --- | --- | --- | --- | --- | --- | --- |
| Task | Effect | $\beta$ | $p$ | $\beta$ | $p$ | $\beta$ | $p$ | $\beta$ | $p$ | $\beta$ | $p$ |
| E1: Logical Syllogisms | Intercept | 0.86 | **0.001** | 2.32 | **<.001** | 1.91 | **<.001** | 1.14 | **<.001** | -0.32 | 0.249 |
|  | Critical | 0.89 | **<.001** | 0.07 | 0.332 | -0.04 | 0.739 | -0.03 | 0.758 | -0.12 | 0.222 |
| E1a: Real-word Syllogisms | Intercept | 0.77 | **0.002** | 2.20 | **<.001** |  |  |  |  |  |  |
|  | Critical | 0.86 | **<.001** | 0.01 | 0.873 |  |  |  |  |  |  |
| E1b: Nonword Syllogisms | Intercept | 0.94 | **<.001** | 2.45 | **<.001** |  |  |  |  |  |  |
|  | Critical | 0.93 | **<.001** | 0.13 | 0.111 |  |  |  |  |  |  |
| E2: Spatial WM MD Loc | Intercept | 0.07 | 0.552 | 1.94 | **<.001** |  |  |  |  |  |  |
|  | Critical | 0.38 | **<.001** | 1.72 | **<.001** |  |  |  |  |  |  |
| E3: Symbolic Syllogisms | Intercept | 1.09 | **0.001** | 2.21 | **<.001** |  |  |  |  |  |  |
|  | Critical | 0.57 | **<.001** | 0.12 | 0.205 |  |  |  |  |  |  |
| E4: Program Induction | Intercept | 0.56 | **0.025** | 1.81 | **<.001** |  |  |  |  |  |  |
|  | Critical | 0.71 | **<.001** | 0.86 | **<.001** |  |  |  |  |  |  |
| E5: Cattell Matrices | Intercept | 0.11 | 0.712 | 2.36 | **<.001** |  |  |  |  |  |  |
|  | Critical | 0.98 | **<.001** | 1.62 | **<.001** |  |  |  |  |  |  |
| E6a: Addition Only MD Loc | Intercept | -0.09 | 0.642 | 0.71 | 0.020 |  |  |  |  |  |  |
|  | Critical | 0.19 | 0.037 | 0.83 | **<.001** |  |  |  |  |  |  |
| E6b: Arithmetic Addition | Intercept | -0.16 | 0.580 | 0.89 | **0.001** |  |  |  |  |  |  |
|  | Critical | 0.03 | 0.812 | 0.63 | **<.001** |  |  |  |  |  |  |
| E6b: Arithmetic Subtraction | Intercept | -0.17 | 0.602 | 1.04 | **0.001** |  |  |  |  |  |  |
|  | Critical | 0.58 | **<.001** | 1.28 | **<.001** |  |  |  |  |  |  |
| E6c: Arithmetic Multiplication | Intercept | 0.44 | 0.099 | 1.50 | **<.001** |  |  |  |  |  |  |
|  | Critical | 0.62 | **<.001** | 1.59 | **<.001** |  |  |  |  |  |  |
| E6c: Arithmetic Division | Intercept | 0.30 | 0.302 | 1.52 | **<.001** |  |  |  |  |  |  |
|  | Critical | 1.25 | **<.001** | 1.70 | **<.001** |  |  |  |  |  |  |
| E7: Language Localizer | Intercept | 0.32 | **0.005** |  |  | 0.73 | **0.007** |  |  |  |  |
|  | Critical | -0.21 | 0.001 |  |  | 1.59 | **<.001** |  |  |  |  |
| E8: Int. Physics Localizer | Intercept | 0.23 | 0.261 |  |  |  |  | 0.23 | 0.385 |  |  |
|  | Critical | 0.17 | 0.046 |  |  |  |  | 0.91 | **<.001** |  |  |
| E9: ToM Localizer | Intercept | 0.54 | 0.112 |  |  |  |  |  |  | -0.16 | 0.589 |
|  | Critical | 0.17 | 0.315 |  |  |  |  |  |  | 1.02 | **<.001** |
| E10: Naturalistic ToM | Intercept | -0.12 | 0.867 |  |  |  |  |  |  | -0.12 | 0.737 |
|  | Critical | 0.56 | 0.216 |  |  |  |  |  |  | 1.27 | **<.001** |
| E10: Naturalistic Social Reln. | Intercept | -0.12 | 0.872 |  |  |  |  |  |  |  |  |
|  | Critical | 0.36 | 0.422 |  |  |  |  |  |  |  |  |

## 1. Logic-sensitive brain regions constitute a network distinct from the Multiple Demand network

Given that the Multiple Demand (MD) network is the leading candidate for domain-general human intelligence (Duncan, 2010, 2013; Duncan et al., 2020), we first examined the response in the MD brain areas to the deductive reasoning task. The MD network was localized with a spatial working memory task where participants have to remember more vs. fewer spatial locations (**Figure 1A**). As expected based on prior research (Fedorenko et al., 2013; Shashidhara et al., 2019; Assem, Blank et al., 2020; Malik-Moraleda et al., 2025), the MD areas responded robustly during this task and showed sensitivity to working-memory load: in left-out data, the hard condition elicited significantly greater activation than the easy condition (*p*<0.001 at the network level; **Figure 1B**, **Table 1**). Critically, in spite of a robust behavioral difference between the hard and easy deduction conditions (accuracies: 91% in the easy condition, 78% in the hard condition, *p*<0.001; RTs: 7.71s in the easy condition, 10.77s in the hard condition, *p*<0.001), the MD network showed little or no response to this contrast, with similarly strong responses to both conditions (**Figure 1B**, **Table 1**). No significant response was present at the network level (*p*>0.3), or in the individual MD regions (**Figure 1B** and **Supp. Table 1**).

Given the surprising result of no response to the Hard>Easy Deduction contrast in the MD network, we looked across the brain for areas sensitive to this contrast. Examination of individual activation maps revealed robust and spatially consistent responses, largely non-overlapping with the MD task activations (**Figure 1C**). This apparent non-overlap was corroborated by the low Dice overlap coefficient (0.02 across the brain and 0.04 for the frontal lobes at the p<0.001 threshold; **Supp. Table 2**; see **Supp. Figure 1** for concordant evidence from voxel-wise spatial correlations).

To formally identify areas of spatially consistent responses across participants, we next performed a whole-brain GSS analysis (Fedorenko et al., 2010; Julian et al., 2012; <u>Methods</u>), which is similar to the traditional random-effects analysis but allows for inter-individual variability in the precise locations of functional areas. This analysis identified 7 regions of consistent activation: 4 in the left frontal lobe (LAF1a,b and LAF2a,b), 2 in the right frontal lobe (RAF1 and RAF2), and 1 in the medial frontal cortex (MF) (**Figure 1D**, <u>Methods</u>). These regions showed, in left-out data, a robust Hard>Easy Deduction effect both at the network level (*p*<0.001; **Figure 1D**, **Table 1**) and in individual regions (*p*s<0.05, after a correction for the number of regions; **Figure 1D; Supp. Table 1**; see **Supp. Figure 3** for additional regions showing weaker and less consistent responses). Furthermore, in contrast to the MD regions, these logic-sensitive regions responded only weakly to the Hard>Easy spatial WM contrast (*p<*0.01 at the network level; note that LAF2a and LAF2b show no hint of this effect). Across the 7 regions, the response to the Hard Deduction condition was more than double the response to the Hard WM condition (**Figure 1D**, **Table 1**), and the logic contrast was reliably greater than in the MD fROIs (*p*<0.001; **Supp. Table 3A**). In contrast, the spatial WM contrast was reliably greater in the MD fROIs (*p*<0.001; **Supp. Table 3B**).

To test whether this dissociation between the logic regions and the MD network holds during naturalistic cognition, we examined the strengths of functional correlations (Yeo, Krienen et al. 2011; Blank et al., 2014; Braga & Buckner, 2017; Du et al., 2024; <u>Methods</u>) during a naturalistic-cognition paradigm (an animated film “Partly Cloudy”; Pixar Animation Studios). The average correlations within each set of regions were high (**Figure 1E**; MD network: 0.48; logic network: 0.44) and higher than the average correlation between the MD and logic regions (0.34; *p*s<0.05).

Finally, a system that supports computations relevant to logical reasoning should be sensitive to problem complexity not only at the coarse condition level, but also at the level of individual problems. Indeed, the level of response in the logic regions to the 32 syllogisms in one of half of the participants could be predicted from the level of response in the other half (R^2^=0.589; *p*<0.0001; **Figure 1F**). In contrast, a similar analysis on the responses in the MD areas to these problems revealed low inter-participant consistency (R^2^=0.106; *p*>0.05).

It therefore appears that deductive reasoning recruits a network of brain areas distinct from the Multiple Demand network.

## 2. Logic brain regions support abstract deductive and inductive reasoning across paradigms

What computations might these logic-sensitive brain areas support? One way to constrain the hypothesis space is by probing these areas’ responses to other reasoning tasks. First, within syllogistic reasoning, we asked whether responses generalize across lexical-semantic content. In Experiment 1, half of the problems used real words (e.g., “If the block is blue, then it is not large…”) and the other half—nonwords (e.g., “If the tep is grix, then it is not ob…”, **Figure 2A**). Because logical structure is content-invariant, logic areas should exhibit the Hard>Easy Deduction effect regardless of the specific labels assigned to the objects and predicates. This is indeed what we found (at the network level: words: *p*<0.001, nonwords: *p*<0.001; **Figure 2B**, **Table 1**). Effect magnitudes did not differ statistically across formats (paired *t-test*, *p*>0.5), and each of the seven logic regions showed significant Hard> Easy effects for both words and nonwords (*p*s<0.05, except LAF1b for nonwords; **Figure 2C; Supp. Table 1**). Next, we tested generalization to non-verbal (symbolic) stimuli, matched for content to the word problems in Experiment 1 (Experiment 3; *n=*14; **Figure 2A**; <u>Methods</u>). The logic network again exhibited a significant Hard>Easy Deduction effect both at the network level (*p*<0.001) and in most individual regions (*p*s<0.05; except LAF1a and RAF1, which responded similarly strongly to both conditions; **Figure 2B–C**, **Table 1; Supp. Table 1**). Thus, the logic regions appear to support deductive reasoning regardless of whether the content of the problem is expressed with words, content-free nonwords, or non-verbal symbols.

**Figure 2.**
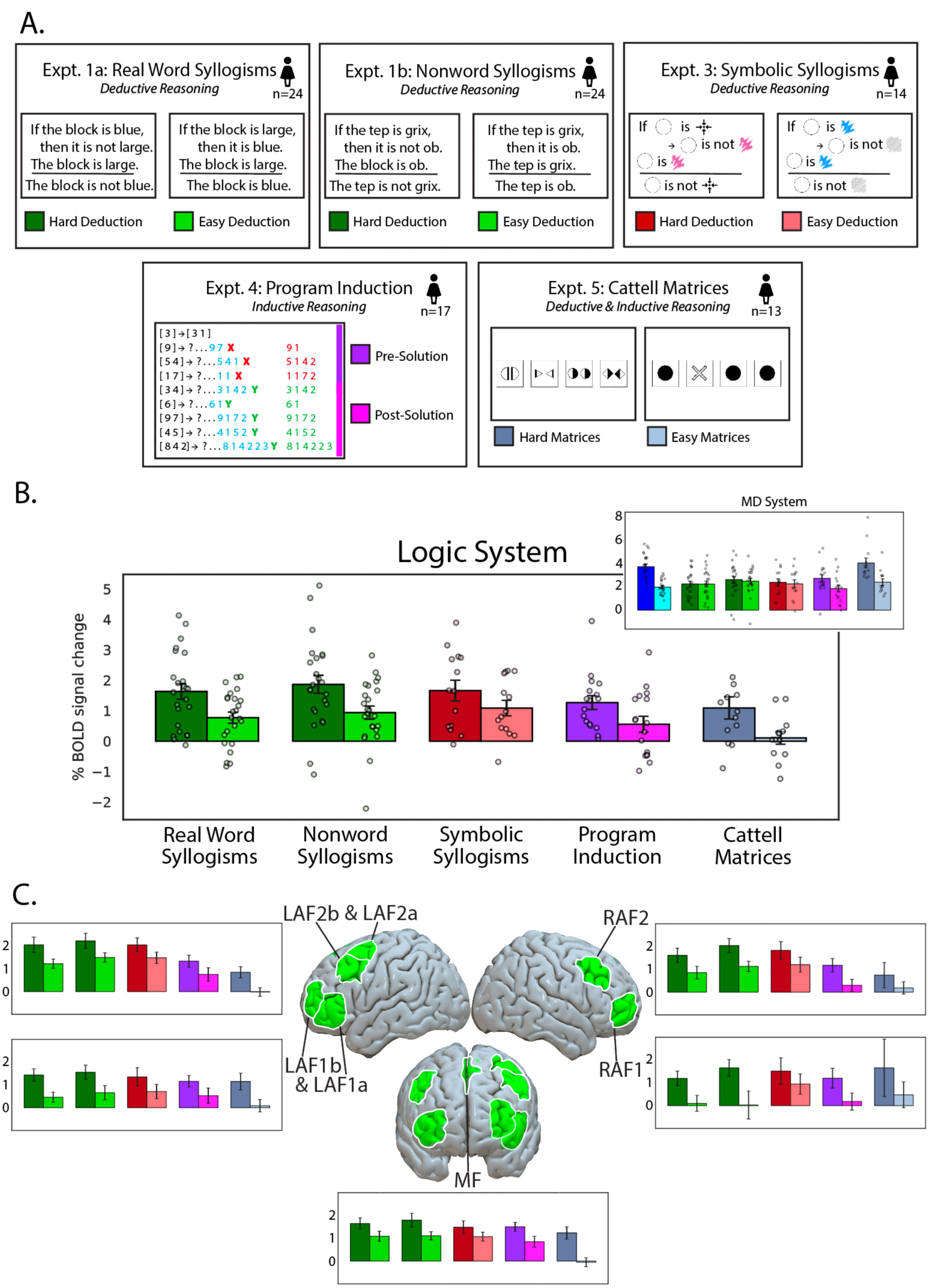
The logic system responds robustly across diverse reasoning paradigms. **A.** Experimental paradigms: Real Word Syllogisms (deductive reasoning, Expt. 1a), Nonword Syllogisms (deductive reasoning, Expt. 1b), Symbolic Syllogisms (deductive reasoning, Expt. 2), Program Induction (inductive reasoning, Expt. 4), and Cattell Matrices (deductive & inductive reasoning, Expt. 5). Each task contrasted Hard vs. Easy conditions. **B.** Responses in the logic system to each paradigm, averaged across fROIs. All five task contrasts elicited reliable Hard>Easy effects. Inset: responses in the MD system to the same tasks, showing little or no selective engagement to the deductive syllogism tasks, but a reliable response to program induction and matrix reasoning. **C.** Responses in individual logic fROIs (LAF1a & LAF1b, LAF2a & LAF2b, MF, RAF1, RAF2). All parcels showed reliable Hard>Easy effects across every reasoning paradigm. Error bars show the standard error of the mean across participants.

We then asked whether the logic regions support another kind of formal reasoning: induction. In contrast to deduction, which derives necessary conclusions from given premises, inductive reasoning involves drawing general principles from specific observations or examples (Peirce, 1878; Pólya, 1954). Whether inductive and deductive reasoning rely on a shared neural substrate remains controversial, with some studies suggesting overlap and others reporting partial dissociations (Goel et al., 1997; Osherson et al., 1998; Parsons & Osherson, 2001; Goel & Dolan, 2004). We adopted a paradigm from Rule et al. (2024; for earlier, related paradigms, see Bruner et al., 1956; Bongard, 1967; Gentner, 1983; Hofstadter & Mitchell, 1994; Tenenbaum, 1999), where participants were presented with an input number list and an output list (e.g., [5, 7] → [7, 5, 7]) and asked to infer the rule that governs the input-to-output transformation. They could then test their hypothesis on a new input list, and so on, until they guess the correct rule. The rules involve a combination of mathematical operations (e.g., *f* (*l*) = [ *x* + 2 ∣ *x* ∈ *l*]), “add 2 to every number”), list operations (e.g., *f* (*l*) = [ *xi* ∣ *xi* ∈ *l*, ∀*j* < *i*, *xi* ≠ *xj*], “leave only unique, non-repeated elements”), and structural operations (e.g., *f* (*l*) = *l*[3:]^*l*[0:3], “rotate the list by three elements”), each of which can be written as short computer programs (Experiment 4; *n=*17; **Figure 2A**; <u>Methods</u>). The logic network showed a reliable response to the contrast between the part of the trial where participants are engaging in inductive reasoning (Pre-Solution) and the part after the correct rule has been inferred (Post-Solution) at the network level (*p*<0.001; **Figure 2B**, **Table 1**) and in most individual regions (*p*s<0.05; except LAF1b; **Figure 2C, Supp. Table 1**), which suggests that the logic regions support not only deductive but also inductive reasoning. The MD network was also engaged by this inductive reasoning task (**Figure 2B inset**; <u>Discussion</u>).

Finally, we examined the logic regions’ responses to a task that was representative of classic IQ-measuring paradigms: an abstract matrix reasoning task (Cattell, 1940; Raven, 2000; Wechsler, 2008). We used an fMRI-adapted version (Woolgar et al., 2013), where participants were shown a matrix of four abstract geometric patterns and asked to identify an outlier. This task engages inductive strategies to generate hypotheses about the rules that govern the given patterns, as well as deductive reasoning to evaluate logical constraints and eliminate incorrect options (Experiment 5; *n*=13; **Figure 2A**; <u>Methods</u>). The logic network showed a reliable response to the Hard>Easy Matrices contrast at the network level (*p*<0.001, **Figure 2B**, **Table 1**) and in most individual regions (*p*s<0.01; except RAF1 and RAF2; **Figure 2C, Supp. Table 1**). As expected based on prior studies (Desco et al., 2011; Woolgar et al., 2013), the MD network also responded to matrix reasoning (**Figure 2B inset, Table 1**).

Together, the data from Experiments 1 and 3-5 suggest that the logic network supports abstract rule and relational reasoning across diverse stimuli (spanning verbal, symbolic, numerical, and visuo-spatial formats) and is engaged by both deductive and inductive inferences.

## 3. Logic brain regions support some aspects of arithmetic reasoning

Next, we investigated the role of the logic network in numerical reasoning. Logic and mathematics are two domains that require formal knowledge representations and have deterministic rules for validity, and thus have often been discussed and studied together in philosophy, cognitive science, and neuroscience (Johnson-Laird, 1983; Rips, 1994; Holyoak & Thagard, 1996; Lakoff & Núñez, 2000; Houdé & Tzourio-Mazoyer, 2003; Goel, 2007; Dehaene, 2008; Holyoak & Morrison, 2012; Monti & Osherson, 2012; Amalric & Dehaene, 2016). Furthermore, according to formal logicist claims, arithmetic is fundamentally grounded in logic (Frege, 1884; Whitehead & Russell, 1910). To investigate whether arithmetic reasoning engages the logic network, we carried out three experiments where participants solved addition problems (Experiment 6a; *n*=10), addition and subtraction problems (Experiment 6b; *n*=14), and multiplication and division problems (Experiment 6c; same *n*=14 as in Expt 6b), with problem difficulty varying within operation type (**Figure 3A**). For the harder multiplication condition, problems were selected so as to minimize the possibility that the answers would be retrieved from rote memorized tables (e.g., 18x7; cf. easier problems whose answers are likely cached, such as 9x7).

**Figure 3.**
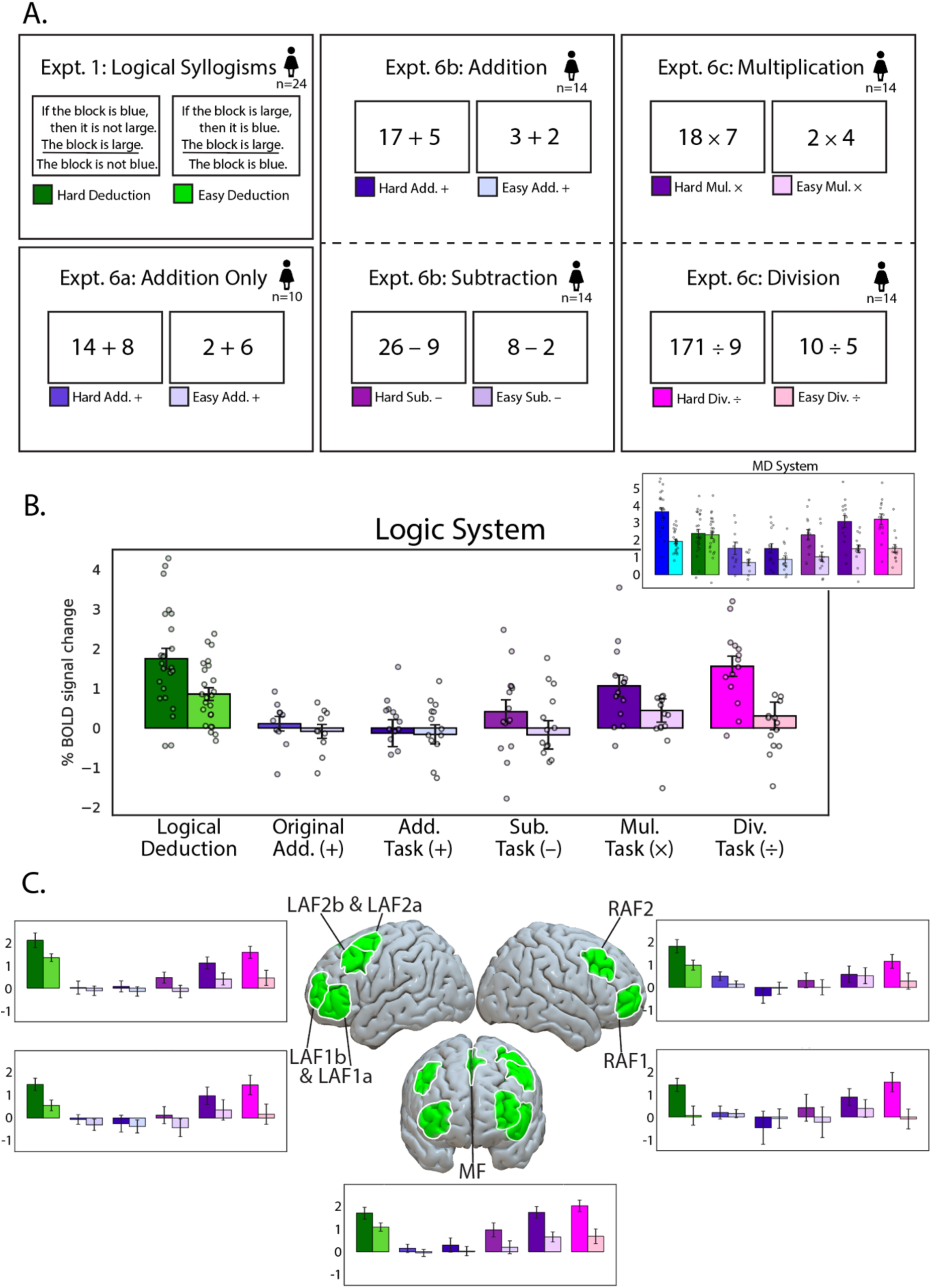
Simple arithmetic engages the MD system but not the logic system, non-cached operations recruit both systems. **A.** Experimental paradigms: participants performed the logical syllogism task (from Expt. 1) and a set of arithmetic reasoning tasks, including addition-only, addition, subtraction, multiplication, and division, each with Hard vs. Easy conditions. **B.** Responses in the logic system across tasks, averaged across fROIs. Logical deduction elicited a robust Hard>Easy effect, but simple addition tasks showed little to no engagement. Inset: responses in the MD system, which showed strong Hard>Easy effects across all arithmetic tasks. **C.** Responses in individual logic fROIs (LAF1a & LAF1b, LAF2a & LAF2b, MF, RAF1, RAF2). All parcels responded robustly to logical deduction, but none showed consistent Hard>Easy effects for simple arithmetic sums.

The logic network showed little to no response to addition problems, with responses close to the fixation baseline in both Experiments 6a and 6b, and no reliable Hard>Easy effect in any fROI (**Figure 3B**, **Table 1**), and only a weak response to the subtraction problems (Hard>Easy effect: *p*<0.001 at the network level; but note that the Hard subtraction condition elicited a response that was substantially lower than the Easy Deduction condition; **Figure 3B**, **Table 1**). The low responses to addition and subtraction were consistent across the logic regions (**Figure 3C; Supp. Table 1**). In contrast, the logic network responded quite strongly to multiplication, and even more strongly to division (Hard>Easy effects: *p*s<0.001 at the network level, and *p*s<0.01 in individual fROIs; **Figure 3B-C**, **Table 1; Supp. Table 1**). The data from these experiments again highlight the dissociation of the logic network from the MD network, which responded strongly and showed robust Hard>Easy effects for all problem types (*p*<u>s</u><0.001; **Figure 3B inset, Table 3**; **Supp. Table 1**).

Together, these results suggest that the logic network is selectively engaged by difficult (non-cached) multiplication and division problems, which, we speculate, require compositional manipulation of relations, such as factor and ratio structure (Discussion).

## 4. The logic network is distinct from the language network and two domain-specific reasoning networks

In Section 1, we showed that the logic network is spatially and functionally dissociable from the Multiple Demand network—the current leading candidate for the neural substrate of fluid intelligence. However, several other brain networks support aspects of high-level cognition and include components in the frontal lobes. Here, we probed the relationship between the logic network and those networks, each identified with an established localizer contrast.

One network that might be expected to overlap with the logic network is the language network (Fedorenko et al., 2010; 2024). Both networks have prominent left frontal components, and language has been argued to provide the structured, compositional format that could support reasoning (Chomsky, 1965; Carruthers, 2002) and perhaps most plausibly, reasoning in sentential or propositional forms of logic. However, past studies found that the language areas do not respond to deductive complexity (Monti et al., 2009; Kean et al., 2025b), and here, we find that the logic network is not engaged during language comprehension (*p>*0.7, **Figure 4B-C**, **Table 1**). These results suggest that the two systems are functionally distinct.

**Figure 4.**
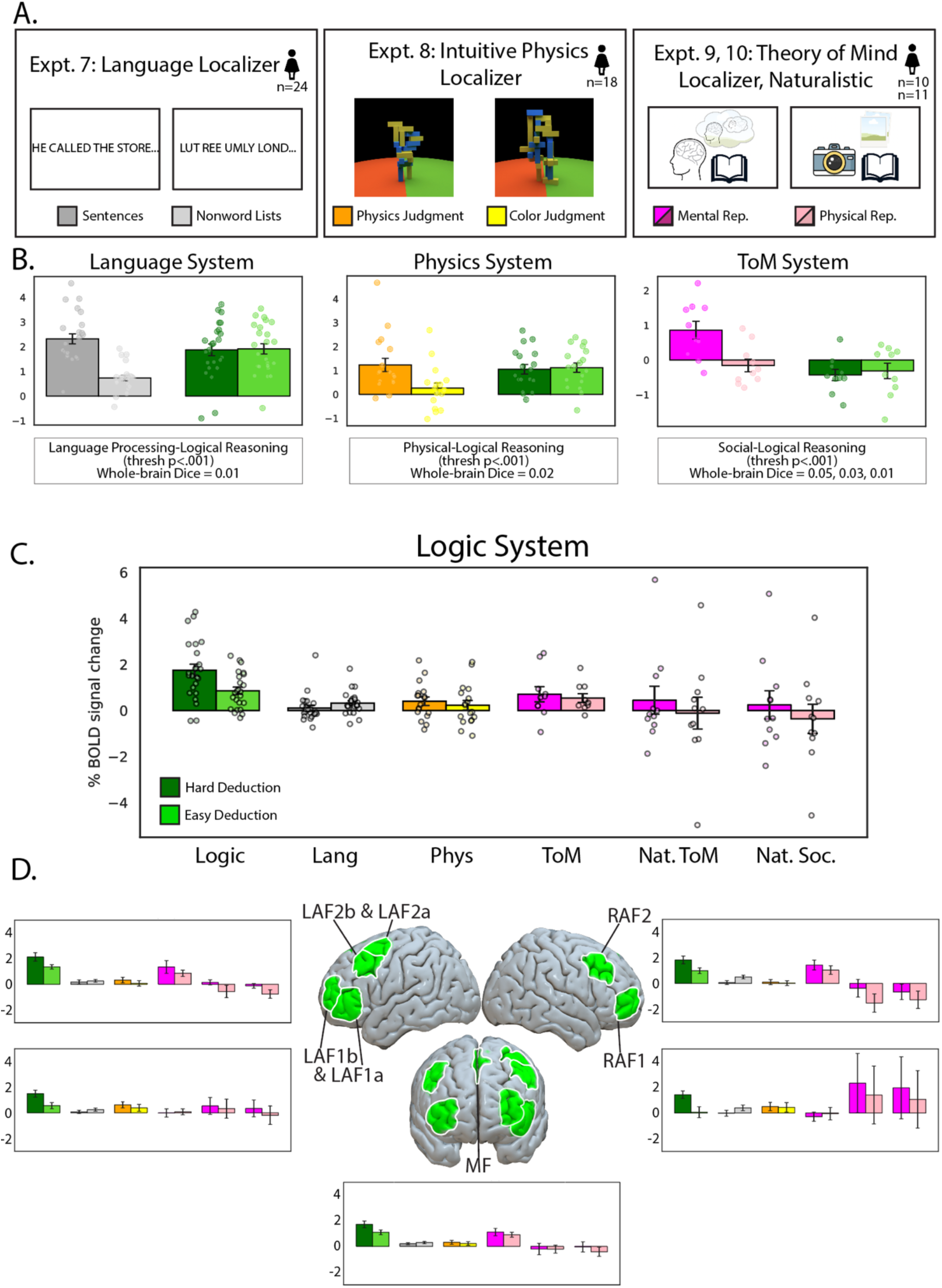
Dissociation between the logic system and linguistic as well as domain-specific reasoning systems (physical and social reasoning). **A.** Experimental paradigms: participants performed a language localizer (Sentences vs. Nonword lists, Expt. 7), an intuitive physics localizer (Physics vs. Color judgments, Expt. 8), a theory of mind localizer (Mental vs. Physical representations, Expt. 9), and a naturalistic social reasoning localizer (Expt. 10) with embedded mental and social contrasts, in addition to the logic localizer (Hard vs. Easy Deduction). **B.** Responses of the language, physical reasoning, and social reasoning systems to their respective native contrasts and to the logic Hard>Easy Deduction contrast. Each system responded robustly to its native contrast but showed no engagement during logical reasoning. Dice coefficients (below) quantify the minimal voxel-wise overlap between each system’s localizer contrast and the logic contrast (p<0.001). **C.** Responses of the logic system to the logic, language, and domain-specific tasks. The logic system showed strong selectivity for logical deduction but no reliable responses to language, physics, or social reasoning contrasts. **D.** Responses in individual logic fROIs (LAF1a & LAF1b, LAF2a & LAF2b, MF, RAF1, RAF2), showing robust Hard>Easy Deduction effects but minimal responses to language and domain-specific reasoning contrasts.

Two other brain networks we considered are domain-specific reasoning systems: the intuitive physical reasoning network, which supports predictive inferences and simulations about objects and events in the world (e.g., rolling, colliding, or falling) (Fischer et al., 2016; Pramod et al., 2022; 2025), and the social reasoning (or Theory of Mind) network, which supports inferences about beliefs, intentions, and other unobservable mental states (Saxe & Kanwisher, 2003; Saxe & Powell, 2006; Castelli et al., 2000; Gallagher et al., 2000; Jacoby et al., 2016). Both networks perform highly abstract computations in their respective domains, suggesting that they could also contribute to more general forms of formal reasoning, and both networks have components in the frontal lobes (Saxe & Kanwisher, 2003; DiNicola et al., 2020; Kean et al., 2025a). However, neither the physical reasoning nor the social reasoning network responded to the logic contrast (**Figure 4B**, **Table 1**), and the responses in the logic network to the physical and social reasoning conditions were overall low (lower than the easy deduction condition) although the physical contrast showed a small but reliable effect (**Table 1**, **Figure 4D; Supp. Table 1**). We complemented these univariate analyses with an examination of fine-grained spatial topographies (**Supp. Figure 1**) and spatial overlap (**Supp. Table 2**), which provided converging evidence for the distinctness of the logic network from these established networks. Together, these results reinforce the logic network as a specialized system for abstract reasoning rather than supporting computations related to linguistic structure, or domain-specific computations related to predictions about the physical or the social worlds.

## Discussion

Using precision fMRI, we have identified a frontal cortical network engaged during abstract formal reasoning. Components of this network can be traced to earlier neuroimaging investigations and neuropsychological patient studies, but here we establish this set of bilateral frontal areas as a functionally integrated system, systematically characterize this network across diverse reasoning paradigms, and establish its distinctness from the Multiple Demand (MD) network and other high-level cognitive networks.

### Dissociation from the Multiple Demand network

Over the last few decades, the Multiple Demand (MD) network has emerged as the leading neural candidate supporting our ability for flexible goal-directed behaviors (Duncan, 2010; Cole & Schneider, 2007; Niendam et al., 2012; Robinson et al., 2022; for earlier theorizing, see Fuster, 1980; Shallice, 1982; Miller & Cohen, 2000). Consisting of bilateral frontal and parietal areas, this network is engaged during diverse cognitively demanding tasks—including the classic suite of executive function tasks—with stronger responses during the more difficult conditions (Duncan & Owen, 2000; Cole et al., 2013; Fedorenko et al., 2013; Shashidhara et al., 2019). This broad engagement suggests that these brain regions may support highly general computations related to attention, working memory (including rule/set maintenance), inhibitory/cognitive control, response selection, and error monitoring (e.g., Duncan, 2010; Fedorenko et al., 2013; Assem et al., 2020). Moreover, based on both brain-behavior correlations in fMRI studies (Tschentscher et al., 2017; Assem et al., 2020) and lesion-symptom mapping (Woolgar, et al., 2010; 2018), the MD network has been implicated in fluid intelligence—a domain-general component of cognition that underlies the *positive manifold* (a positive correlation in performance across tasks; Spearman, 1904, 1927; Carroll, 1993; Thorndike, 1994; Jensen, 1998; van der Maas et al., 2006; Kovacs & Conway, 2016). Logical reasoning certainly constitutes a case of demanding cognition, which requires attention, places demands on working memory, and so on. Indeed, the logical reasoning task elicited a strong response in the MD network. Critically, however, the MD areas showed no sensitivity to logical complexity. Instead, a set of distinct areas showed such sensitivity, scaling with logical complexity manipulations, but showed no response to a demanding spatial working memory task, which robustly engages in the MD network. This dissociation suggests that logical reasoning is not reducible to the kind of cognitive demands supported by the MD network.

Some proposals of the organization of the prefrontal cortex (PFC) posit a rostro–caudal gradient of increasingly abstract cognitive control demands toward the anterior-most portions of the PFC (Koechlin et al., 1999; 2003; Koechlin & Summerfield, 2007; Badre & D’Esposito, 2007; Badre & Wagner, 2007; Badre, 2008; Nee & D’Esposito, 2016; Thiebaut de Schotten et al., 2017). This idea has been difficult to reconcile with the existence of the MD network, which appears to be recruited as a whole, including its frontal components, even during relatively simple task demands (Crittenden & Duncan, 2014). Our data may help reconcile these two lines of evidence: instead of the rostro-caudal gradient, prior findings may be explained by the existence of two nearby brain networks: the logic network has a frontal component that lies anteriorly to the lateral frontal MD areas and may implement more abstract and complex forms of behavioral control than those supported by the MD network (see also Badre & Nee, 2018).

Does the logic network represent the apex of cognitive control—a hierarchical endpoint in the same ‘lineage’ as cognitive control, where regulation reaches its most abstract and relational form? Or does it constitute an independent faculty with a distinct line of descent, a parallel system that follows a separate developmental and evolutionary trajectory while interfacing with control when needed? These alternatives—to be distinguished in future work—imply very different architectures of thought: the first casts human reasoning as the continuing arc of domain-general control, but the second points to a qualitatively distinct, and possibly uniquely human, cognitive capacity.

The frontal lobe locus of the logic network is notable. The frontal lobes have long been regarded as the seat of higher cognition and goal-directed behavior (Harlow, 1848; 1868; Miller & Cohen 2000). With the discovery of the MD network, which has a prominent parietal component—along with the fact that the human association cortex expanded across not only frontal, but also parietal and temporal lobes (Buckner & Krienen, 2013)—the focus has gradually shifted away from the anterior portions of the brain. Although the logic network does have parietal components (**SI-3**), those components are less robust and less spatially consistent across individuals. Thus, the privileged role of the frontal lobes in human cognition may need to be revisited.

### Formal vs. domain-specific reasoning

In addition to the MD network, two other networks have been implicated in abstract reasoning, but for particular content domains: social reasoning and reasoning about the physical world. The social reasoning (or Theory-of-Mind) network includes components in the bilateral temporo-parietal junction and along the cortical midline (Fletcher et al., 1995; Castelli et al., 2000; Gallagher et al., 2000; Saxe & Kanwisher, 2003; Ruby & Decety, 2003; Saxe, et al., 2006), and the physical reasoning network comprises lateral frontal and lateral parietal areas bilaterally (Fischer et al., 2016; Schwettman et al., 2019; Pramod et al., 2022; 2025). A critical shared feature of these two domain-specific reasoning networks is the abstractness of the representations they handle. Both networks process the relevant content across diverse input formats (Koster-Hale et al., 2017; Castelli et al., 2000; Gallagher et al., 2000; Jacoby et al., 2016; Pramod et al., 2022; 2025), suggesting that they represent mental state information or physical relationships abstractly.

This representational abstractness of social and physical reasoning has sometimes led researchers to treat these forms of reasoning as proxies for domain-general logical thought. Social intelligence in dolphins and primates, for example, has been argued to rely on knowledge of the hierarchical rules governing alliances, dominance, and cooperation (Byrne & Bates, 2007; Connor, 2007). Likewise, complex forms of tool use in primates suggest sensitivity to abstract causal and relational structures beyond the immediate perceptual input (Seed et al., 2011; Gärdenfors & Lombard, 2020). Furthermore, our lived experience is fundamentally grounded in the physics of our world, and in the social structures within which we exist (especially for primates; Cheney et al., 1986; Seyfarth & Cheney, 2015). We begin learning about the physical and social worlds from the first days and weeks of our lives, and some have even hypothesized that aspects of physical and social knowledge may be innate (Baillargeon, 1994; Hespos & Baillargeon, 2001; Baillargeon, 2004; Spelke & Kinzler, 2010). However, in spite of the importance of abstract physical and social reasoning in human lives, humans are also able to reason about elements that are only linked to one another, without being grounded in the external world. This kind of reasoning is harder and more error prone (Wason, 1966, 1968), and grounding an abstract problem within a social context facilitates reasoning (Griggs & Cox, 1983; Cosmides & Tooby, 1992; Evans & Clibbens, 1995), but humans still can and do engage in reasoning about fully abstract structures and relationships (Holyoak & Morrison, 2012; Harel & Soto, 2016). And the newly identified logic network is likely a critical substrate for such reasoning.

Although some have proposed that language—which is by its nature abstract and structured—may endow humans with the ability for abstract reasoning (Vygotsky, 1934; Chomsky 1993; 1995; Carruthers, 2002; Gentner, 2016; Spelke, 2017), ample evidence suggests that the nature of representations mediating thought are non-linguistic, including arithmetic and mathematical reasoning (Varley et al., 2005; Fedorenko et al., 2011; Monti et al., 2012), computational/programmatic reasoning (Ivanova et al., 2020; Liu et al., 2020), social reasoning (Varley & Siegal, 2000; Apperley et al., 2004; 2006; Paunov et al., 2022; Shain et al., 2023), physical reasoning (Kean et al., 2025a), and causal reasoning (see Fedorenko et al. 2024 for a review). In line with this past evidence, we also show that the logic network is distinct from the language network (see also Kean et al. 2025b).

### What computations does the logic network support?

Given that the logic network responds to both deductive and inductive reasoning and appears to contribute to solving non-cached arithmetic problems, which require multi-step reasoning about factor and ratio structure, the most parsimonious account of these brain regions is that they represent and process structured relational information, compute the specific constraints that license particular inferences, and update mental structures after inferential transformations. The precise nature of the computations underlying logical inference remains debated, and our findings can inform these debates. For example, according to the *mental model* (or *simulation*) framework, reasoning entails constructing iconic, structure-preserving analogs of problems, and is supported by spatial or domain-general working memory resources (Johnson-Laird, 1998; 2010; Knauff, 2013; cf. Khemlani & Johnson-Laird, 2013 for a proposal of more abstract representations). Our findings suggest that such resources are not sufficient for logical reasoning: the Multiple Demand network, which supports domain-general working memory, is active during logical reasoning but is not sensitive to logical complexity. Similarly, approaches that emphasize *schemas or scripts*—domain-specific relational templates that scaffold reasoning with content-rich information (Piaget, 1923; Bartlett, 1932; Rumelhart, 1975)—are unlikely to explain our findings, given that our deductive and inductive reasoning paradigms engage brain areas distinct from the domain-specific reasoning systems.

Two classes of accounts are more easily compatible with the existence of brain regions that support abstract logical reasoning. First, classic *mental logic* theories and some variants of the *Language of Thought* (LOT) hypothesis treat reasoning as the symbolic manipulation of abstract propositions according to inference rules (Rips, 1983; 1994; 1995; Fodor, 1975; Mandelbaum et al., 2022; Quilty-Dunn et al., 2022; 2023). Probabilistic extensions of LOT (Tenenbaum et al., 2011; Goodman & Stuhlmüller, 2014; Ellis, 2020; Rule et al., 2020) invoke program-like operations over abstract concepts, encoding both probabilistic knowledge and relational structure. Our findings are consistent with these symbolic accounts, postulating amodal representations that encode inference-relevant structure, although our data add to prior evidence against linguistic variants of the (P)LOT and mental logic theories (Shea, 2024). Finally, *pragmatic or socially-grounded theories* (Cheng & Holyoak, 1985; Cosmides, 1989; Cosmides & Tooby, 1992; Gigerenzer et al., 1999; Lombrozo, 2006; Mercier & Sperber, 2011) remain broadly compatible with our findings, insofar as they allow for a dedicated mechanism for evaluating reasons or logical forms with respect to their validity, independent of any specifically social or pragmatic context.

### A candidate for a human-specific adaptation?

On the archaeological record, the capacity for explicit, domain-general, formal logic is a remarkably late cultural achievement. It emerges substantially later than other cultural advances, including language, which was plausibly in place as early as a million years ago (Dediu & Levinson, 2013), complex hierarchical toolmaking, such as Levallois core technology, which became widespread after ∼300–250k BCE (earliest claims ∼400 k BCE; Bar-Yosef & Van Peer, 2009; Shea, 2013), and symbolic tallies for quantification, which appeared by ∼43–42k BCE (Lebombo bone; d’Errico et al., 2012). Even the earliest written if–then corpora, such as the Ur-Nammu law code (∼2100 BCE), remain narrowly domain-bound (Roth, 1995). Transferable, content-independent procedures emerge only in Old Babylonian and Egyptian mathematics (∼2000–1600 BCE), including completing the square, approximating √2, and the method of false position (Høyrup, 2002; Robson, 2008; Friberg, 2007; Clagett, 1999; Imhausen, 2016), culminating in the fully formal, domain-general systems of the first millennium BCE (Euclid, Aristotle, Nyāya, Mohist Canons; Matilal, 1998; Ganeri, 2001; Graham, 1978; Fraser, 2016).

Given the late cultural emergence of domain-general logical computation, this system may be uniquely human. The Multiple Demand system appears to be broadly conserved across primates (Mitchell et al., 2016), but nonhuman primates show limited generalization of formal rules without extensive training (Cantlon et al., 2024; Goudar et al., 2024). One possibility then is that at some point during the evolutionary expansion of the frontal lobes (Buckner & Krienen 2013), a logic-selective network may have fractionated from the MD system and eventually assumed its current functional form as cultural practices demanded fully explicit, content-independent reasoning. Indeed, the frontal polar areas—which house the anterior-most components of the logic network (areas LAF1 and RAF1)—have been identified as exhibiting substantial differences between humans and non-human primates in size, cytoarchitecture, and functional connectivity (Semendeferi et al., 2001; Neubert et al., 2014; Donahue et al., 2018; Bryant et al., 2025), suggesting they have undergone substantial reorganization in humans. Studying abstract reasoning in NHPs is far from trivial (Penn et al., 2008; Vendetti & Bunge, 2014; Schubiger et al., 2020; Barron et al., 2020), but modern approaches to cellular-level cross-species comparisons (Suresh et al., 2023; Zemke et al., 2023) may eventually tell us whether there exist true homologies to the logic areas in non-human primate brains.

### Limitations and open questions

We have only sampled some forms of reasoning in one population—neurologically healthy educated adults from an industrialized culture (Henrich et al., 2010). Is the logic system we have identified ubiquitous across humans? Would other forms of reasoning, such as analogical and relational reasoning, probabilistic reasoning, counterfactual reasoning, and reasoning in contextualized/grounded paradigms recruit the logic system? How does the logic system divide labor with the Multiple Demand system and with other reasoning systems? How does the logic system develop, and is it similar in expert logicians and mathematicians? Given the dissociation between the logic system and the MD system, can we find double dissociations in behavior, in patients with brain damage or in investigations of individual differences? Are the different regions within the logic network functionally similar, or do they contribute in distinct ways to logical reasoning? All these questions can and will be answered with additional neuroscientific and behavioral investigations.

Finally, understanding the precise computations carried out by the logic regions—and other areas supporting high-level cognition—remains a lofty goal. Here, of most promise are deep imaging approaches, where brain responses are obtained to hundreds of individual reasoning problems/problem-types. Such data (currently only available for perceptual domains; Chang et al., 2019; Allen et al. 2022; Herbart et al., 2023), in combination with direct comparisons between human neural data and computational models (Yamins & DiCarlo, 2016; Schrimpf et al., 2020), may bring us closer to understanding the algorithms of human thought.

## Methods

### Participants

Twenty-four participants contributed data to the fMRI component of the study (14 female; mean age = 26.4 years, SD = 8.6); all but two participants were right-handed; the one ambidextrous and one left-handed participants had left-lateralized language systems, as determined by the language localizer task described below. The participants were recruited from MIT and the surrounding Cambridge/Boston, MA, community and paid for their participation. All were native English speakers, had normal hearing and vision, and had no history of language or cognitive impairment. All participants provided written informed consent in line with the requirements of MIT’s Committee on the Use of Humans as Experimental Subjects.

All participants completed the logical syllogisms reasoning task, the language localizer, and the Multiple Demand (MD) localizer (spatial working memory task). Different subsets of the 24 participants completed the other tasks, with at least 10 participants performing each task (14 completed the symbolic syllogisms task; 17 completed the inductive reasoning task; 13 completed the matrix reasoning task; 10 completed the addition-only arithmetic task; 14 completed the addition & subtraction task; 14 completed the multiplication & division task; 18 completed the intuitive physical reasoning localizer; 11 completed the theory of mind (ToM) localizer; and 11 completed the alternative ToM localizer based on a naturalistic animated film).

### Reasoning Tasks

The materials for all tasks are available at OSF (osf.io/hqa23).

#### Verbal deductive syllogisms task

Participants were presented with classic three-sentence syllogisms and asked to judge the validity of the third sentence (the conclusion) given the first two sentences (the premises), by pressing one of two buttons on a button box. Each trial corresponded to a syllogism, and trials varied in difficulty between harder deduction (Modus Tollens) and easier deduction (Modus Ponens), and in whether the problems used real words or nonwords; the experiment also included two conditions of no interest (challenging memory conditions). The Modus Tollens > Modus Ponens (Hard Deduction > Easy Deduction) contrast targets cognitive processes related to the complexity of logical deductive reasoning. The trials were distributed across four scanning runs, with 8 trials per condition per run and each run lasting between 551 s and 1,037 s (durations are variable given the self-paced nature of the task). Each participant completed between 2 and 4 runs (16-32 trials per condition), with the condition order counterbalanced across runs (here and in all other experiments).

#### Symbolic deductive syllogisms task

This task was identical to the verbal syllogisms task, except that the nouns and adjectives were replaced with symbolic pictograms. The trials were distributed across two scanning runs, with 8 trials per condition per run and each run lasting between 167 s and 451 s (durations are variable given the self-paced nature of the task). Each participant completed two runs (16 trials per condition).

#### Program induction task

Participants were shown an input list of 1-5 single-digit numbers and an output list of 0-5 single-digit numbers and asked to guess the rule that transformed the input list into the output list. They were then shown another input list and asked to provide the output list. They were told whether or not their guess about the underlying rule is correct and are shown another input list. The responses were entered via a scanner-safe fiber-optic button response device (Nata Technologies LxPad system), which contains 12 buttons allowing input of digits 0–9, backspace, and enter. Each trial (corresponding to a rule) consisted of 8 problems. After the eighth problem, participants were told—via a schematic (**Fig. 2A**)—the correct rule and were asked to apply this rule to two more input lists. Because in most cases, participants guessed the correct rule part-way through the initial list of 8 problems (as indicated by generating correct output lists for the last several problems), we defined the *induction* period as the subset of the eight problems before the problem after which the participant made no more errors, and the *application* period as the subsequent problems along with the two problems after the correct rule was revealed. The induction > application contrast targets neural processes involved in hypothesis generation and rule discovery beyond those required for rule implementation and response entry. Each participant completed 40 rules. The rules were distributed across 20 scanning runs, with 2 rules per run and each run lasting between 200s and 420s (durations are variable given the self-paced nature of the task).

#### Matrix reasoning

Participants were presented with sets of four images of geometrical shapes and asked to decide which image does not fit with the others, by pressing one of four buttons on a button box. The experiment used a blocked design; trials in the hard blocks used stimuli from Cattell (1940), and trials in the easy blocks used simpler problems created by Woolgar et al. (2013), who adapted this task for fMRI (e.g., three images of the same simple shape and an image of a different shape). The hard > easy contrast targets cognitive processes related to relational reasoning, hypothesis generation, and deductive inference. Each participant completed a single run of the task lasting 320 s and consisting of 8 easy blocks and 8 hard blocks. The blocks were of fixed length (16 s), which means that only a few hard trials could be solved during this period (between 1 and 6), and more easy trials could be solved (between 4 and 15); furthermore, because only 24 hard items were available, some participants did not have enough items for all 8 blocks, but every participant completed at least 5 hard blocks.

#### Arithmetic reasoning task (Addition; Addition & Subtraction)

Participants had to solve addition problems (Experiment 6a) or addition & subtraction problems (Experiment 6b) with smaller (easy condition) versus larger (hard condition) numbers, in a blocked design. In the easy condition, participants added two single-digit numbers, or subtracted one single-digit number from another single-digit number. In the hard condition, participants either added two numbers, one of which was double-digits or subtracted a single-digit number from a double-digit number. In both conditions, participants performed a two-alternative forced-choice task at the end of each trial to indicate the correct answer. Each trial lasted 3 s. Each block consisted of five trials and lasted 15 s. Each run consisted of 16 experimental blocks (eight per condition) and five fixation blocks (15 s in duration each), for a total duration of 315 s (5 min, 15 s). All participants performed two runs.

#### Arithmetic reasoning task (Multiplication & Division)

Participants had to solve multiplication & division problems with smaller (easy condition) versus larger (hard condition) numbers, in a blocked design. In the easy condition, participants either multiplied single-digit with single-digit numbers (or 2 times 12-19, or 3 times 12-15) or solved the inverse division problems. In the hard condition, participants either multiplied a single-digit number (excluding 2 and 3) with a double-digit number (excluding some easier multiples of 5), one of which was double-digits, or they solved the inverse division problems. In both conditions, participants performed a two-alternative forced-choice task at the end of each trial to indicate the correct answer. Each trial lasted 4 s. Each block consisted of five trials and lasted 20 s. Each run consisted of 16 experimental blocks (eight per condition) and five fixation blocks (15 s in duration each), for a total duration of 395 s (6 min, 35 s). All participants performed two runs.

For the descriptions of the canonical localizer tasks for the Multiple Demand, Language, Intuitive Physics, and Theory of Mind systems (all used in numerous prior published studies), see **Supp. Methods**.

#### fROI definition and response estimation

(fMRI data acquisition, preprocessing, and modeling, described in **Supp. Methods.**)

#### Definition of the deductive logical reasoning fROIs

We first performed a group-constrained subject-specific (GSS) analysis (Fedorenko et al., 2010; Julian et al., 2012) on the data from Experiment 1 in order to create a set of parcels that would be used for defining individual-level fROIs. GSS is a whole-brain analysis that identifies spatially consistent (across participants) areas of activation for some contrast of interest. To do so, we first thresholded the individual t-maps for the Hard > Easy Deduction contrast by selecting the 10% of most responsive voxels across the brain. These maps were then binarized (selected voxels were turned into 1s and the remaining voxels into 0s) and overlaid to create a probabilistic overlap map (summing the ones and zeros across participants in each voxel). After dividing the summed value in each voxel by the number of participants (n=24 in this case), these values can be interpreted as the proportion of the participants for whom that voxel belonged to the top 10% of most responsive voxels. This probabilistic overlap map was then thresholded, such that voxels with values of 0.1 or lower were removed, and a watershed algorithm was used to segment the map into discrete regions (parcels).

The resulting 25 parcels were evaluated on two criteria. First, we tested whether the Hard > Easy Deduction contrast was replicable across runs. To do so, we used odd-numbered runs to define the functional regions of interest (fROIs) (as the top 10% of most responsive voxels within each parcel, based on the t-values for the Hard > Easy Deduction contrast) and even-numbered runs to estimate the responses; then we did the opposite; and finally, we averaged these estimates to obtain a single estimate per participant per parcel. The Hard > Easy Deduction contrast reliably differed from zero in 21 of the 25 regions. Next, we examined the proportion of participants showing a non-zero intersection with each parcel (i.e., having at least one supra-threshold voxel within the parcel boundaries). The seven parcels where 23 of the 24 or all 24 participants (>95%) had task-responsive voxels were included in the main analyses (**Fig. 1D**; see **Supp. Fig 3** for the visualization of the lower-overlap parcels). To define the individual fROIs, we selected the top 10% of voxels within each of these 7 parcels based on the individual Hard > Easy Deduction activation maps.

fROI definition for the Multiple Demand, Language, Intuitive Physics, and Theory of Mind systems is described in **Supp. Methods.**

### Critical Analyses

In all analyses, the regions of interest were defined functionally in each individual participant (as the top 10% of most responsive voxels for the relevant contrast within each parcel/mask; for the logic network, the parcels were created in the current study as described in the main text; for all other networks, the parcels were created in past work from activation maps from large numbers of participants). To estimate the responses in a region to the conditions used to functionally define the region, a split-half approach was used to ensure the independence between the data used for fROI definition vs. response estimation (e.g., Kriegeskorte, 2011). The responses in each fROI and set of fROIs (for network-level analyses) were statistically evaluated using linear mixed-effects models implemented in R (lme4 package; Bates et al., 2015), with random intercepts for participants and regions. P-values were approximated using the lmerTest package (Kuznetsova et al., 2017), and effect sizes (Cohen’s *d*) were estimated using the EMAtools package (Kleiman, 2017). For each contrast, we fit separate models for the network and for individual fROIs:

The **network-level model**: *BOLD ∼Condition + (1 | Participant) + (1 | fROI)*.

The **individual fROI-level model**: *BOLD ∼Condition + (1 | Participant)*.

Functional connectivity, spatial correlation, and dice overlap analyses are described in **Supp. Methods**.

## Supplementary Information

**Supplementary Figure 1.**
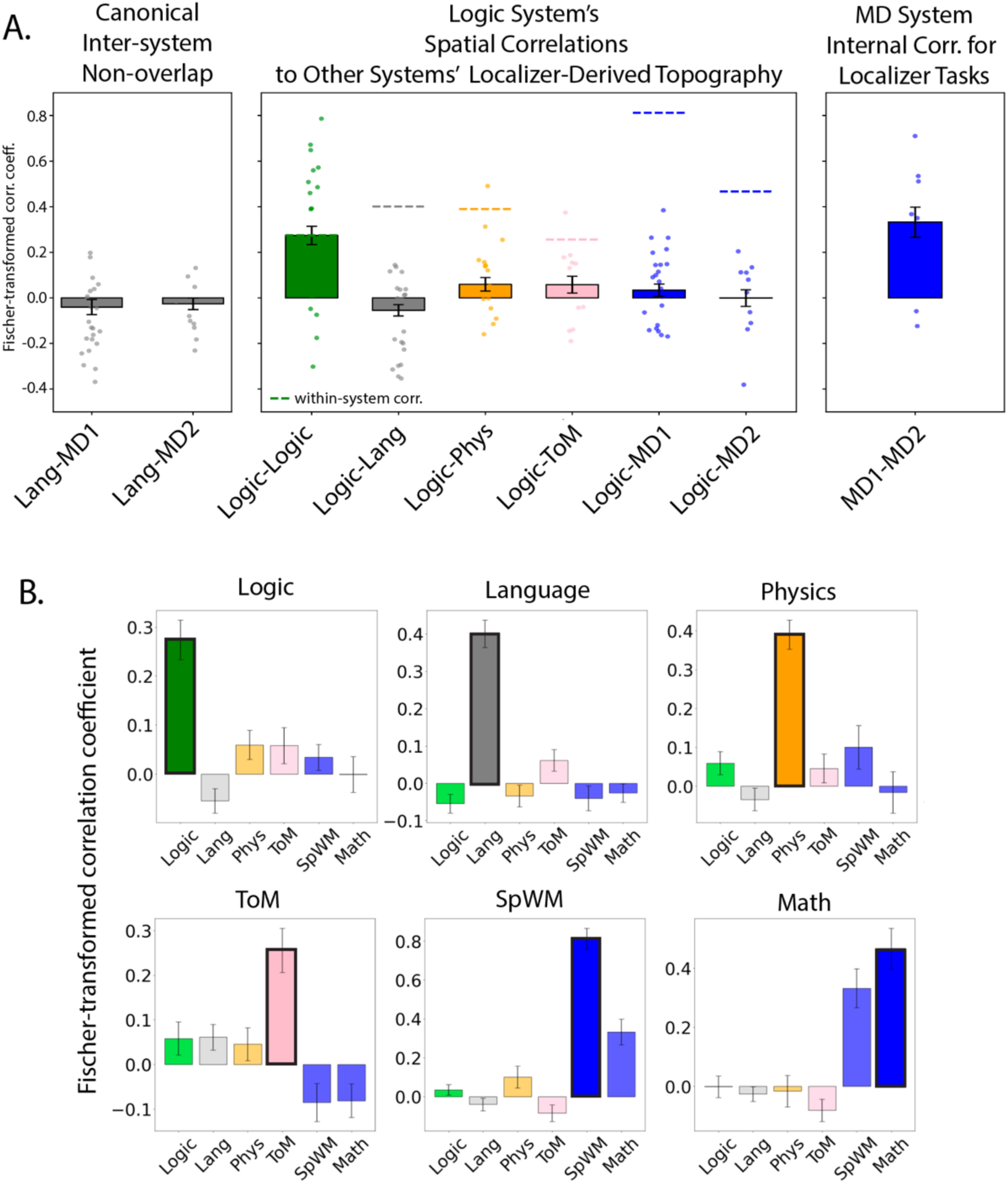
Within- and between-system spatial correlations. **A.** Middle panel: The logic system’s correlation with itself (green; computed using a split-half approach, between odd and even runs of the fMRI task) and with other systems (grey: language; orange: intuitive physics; pink: ToM; and blue: MD – the two bars correspond to the spatial WM task and the arithmetic addition task, both commonly used as the MD system localizers). The dashed line above the non-logic system bars show the within-system correlations, which set the upper bound for the between-system correlations. For comparison, we show correlations between systems known to not overlap: language and MD (Left panel), and between tasks known to engage the same, MD, system: spatial working memory and arithmetic (Right panel). **B.** Each non-logic system’s correlation with itself (darker bars with a black outline in each panel) and with each of the other systems (lighter bars; colors same as in A). For all systems, the within-system correlation is high, whereas the between-system correlations cluster around zero.

**Supplementary Figure 2.**
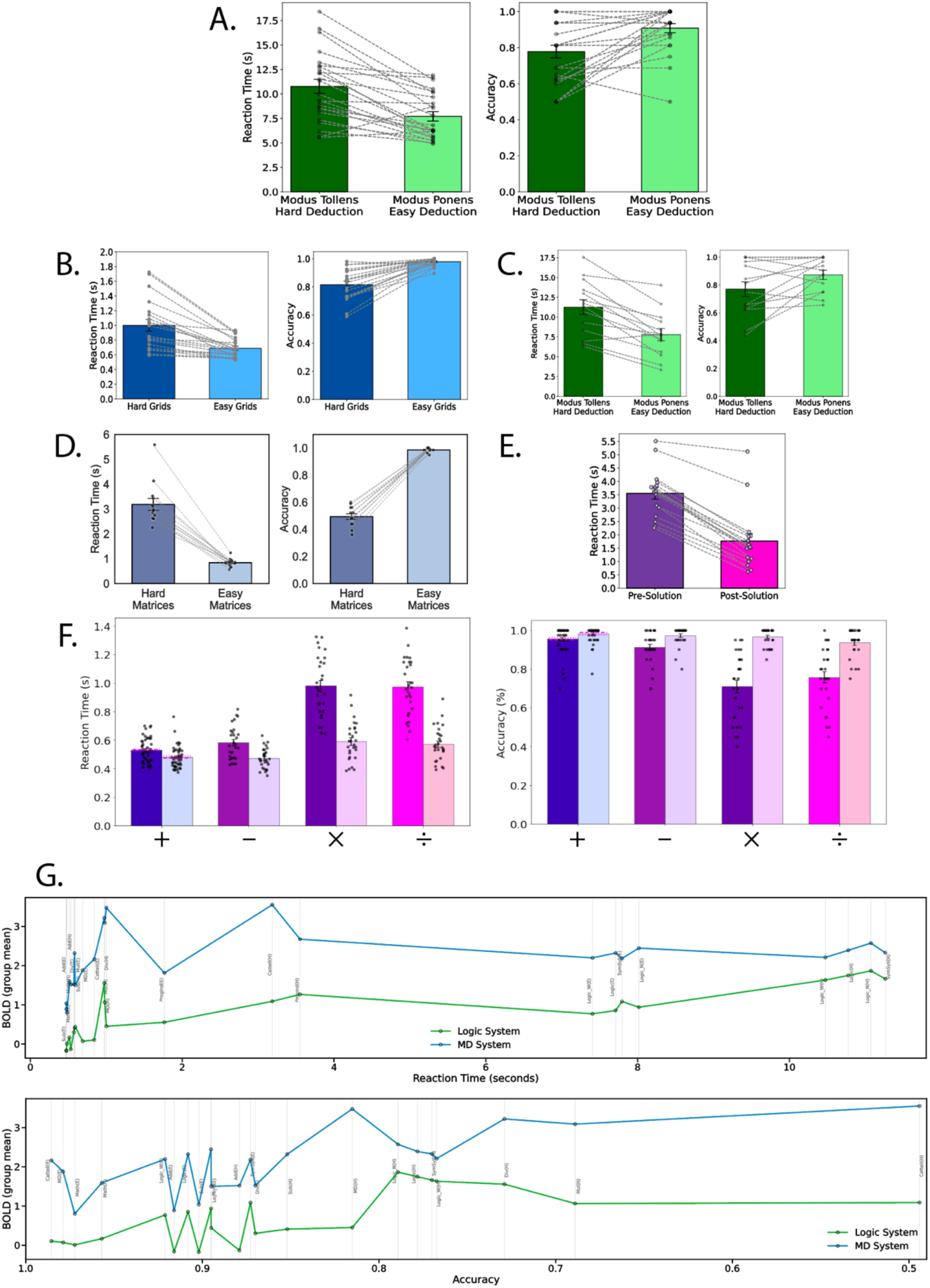
Reaction Times and Accuracies in the Verbal deductive syllogisms task, spatial working memory task, and other reasoning-related tasks. Each panel shows mean reaction times (left) and accuracies (right), with individual participants connected by dashed lines. **A.** Verbal deductive syllogisms (Experiment 1): dark and light green bars represent *Hard Deduction* (*Modus Tollens*) and *Easy Deduction* (*Modus Ponens*) conditions, respectively. Participants were more accurate (*t* = 3.775, *p* < 0.001) and faster (*t* = –5.641, *p* < 0.001) in the easy (*Modus Ponens*) condition. **B.** Spatial working-memory task (Experiment 2): dark and light blue bars correspond to the *Hard* and *Easy* spatial working memory grid configurations, showing the expected effect of task difficulty. **C.** Symbolic syllogisms (Experiment 3): dark and light green bars again represent *Hard Deduction* (*Modus Tollens*) and *Easy Deduction* (*Modus Ponens*) conditions, replicating the verbal syllogisms’ pattern with symbolic stimuli. **D.** Program induction (Experiment 4): dark purple and light pink bars correspond to *Pre-Solution* and *Post-Solution* phases, showing faster responses once the underlying rule has been discovered. **E.** Cattell matrices (Experiment 5): dark and light blue bars indicate *Hard* and *Easy* matrices, with easier items producing faster and more accurate performance. **F.** Arithmetic reasoning (Experiments 6a–c): bars from left to right represent *Addition (+)*, *Subtraction (–)*, *Multiplication (×)*, and *Division (÷)*, from blue, to purple, to violet, to pink, lighter indicates *Easier operations* and darker indicates *Harder operations*. As in other tasks, harder operations elicited longer reaction times and lower accuracies. For the addition condition, pink dashed and dotted lines show group means from Experiments 6a and 6b, respectively, while the bar reflects the overall mean across all addition problems. **G.** Responses of the Logic system (green) and Multiple-Demand (MD) system (blue) are shown across all task conditions as a function of task difficulty, with reaction time on top and accuracy on the bottom.

**Supplementary Figure 3.**
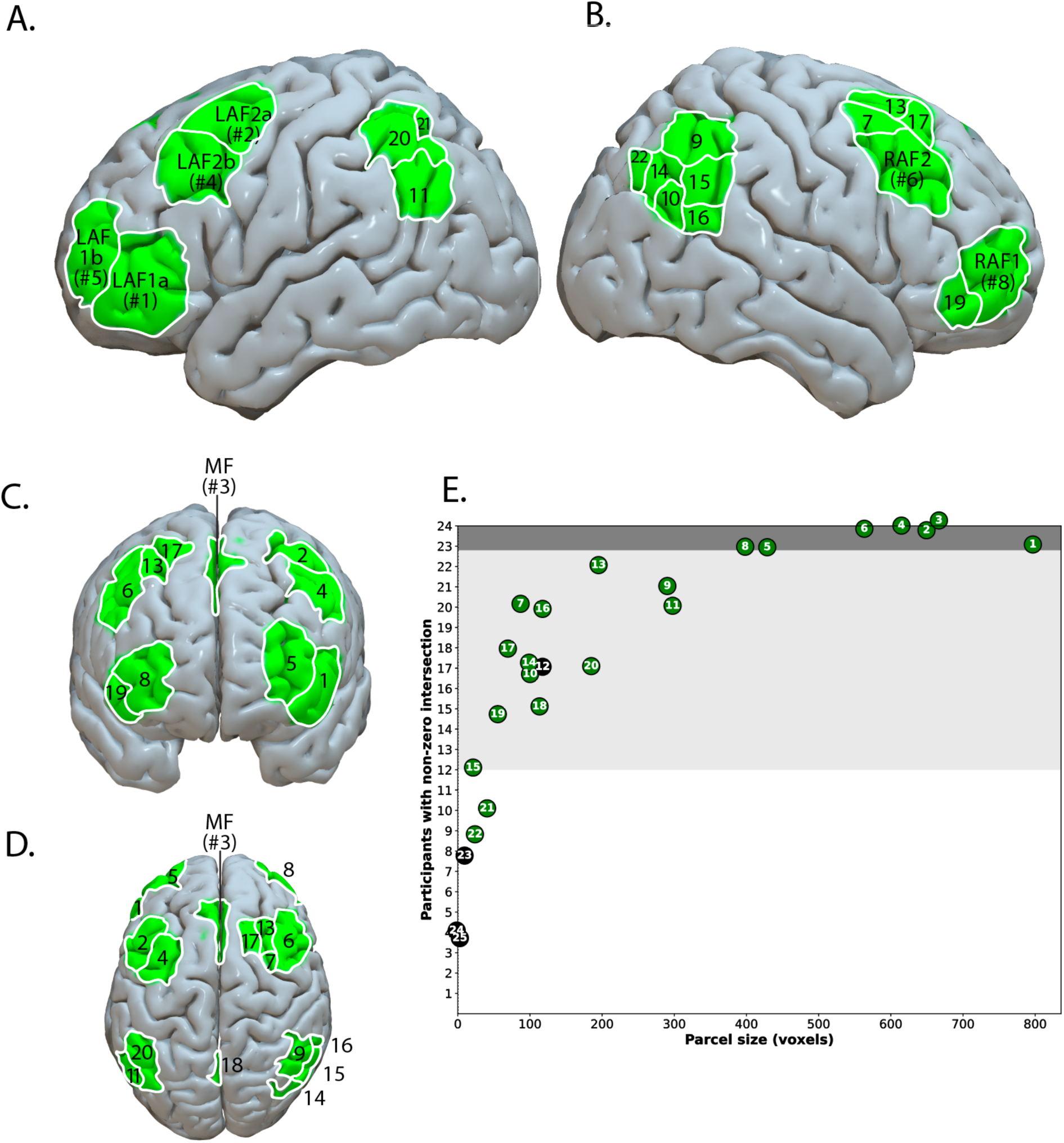
The full set of Logic parcels derived from a whole-brain group-constrained Subject-Specific (GSS) analysis. A–D. Left lateral, right lateral, anterior, and superior views of parcels. These visualizations include the 21 parcels that resulted from the GSS analysis and passed our selection criteria, i.e., showing a reliable Hard > Easy Deduction effect in a left out run of the data. (see <u>Methods</u>). E. The relationship between the size of the parcels (x-axis) and the number of participants that have a non-zero intersection with that parcel (i.e., at least 1 voxel within the borders of the parcel was selected in that participant) (y-axis). The parcels (n = 7) that have a non-zero intersection with all (or 23 of the 24) participants and were selected as the main set for all analyses fall within the dark gray area; the parcels that have a non-zero intersection with 12 or more (≥50%) of the participants fall within the lighter gray area; and the remaining parcels (present in fewer than half of the participants) fall within the white area. Furthermore, the parcels indicated with green dots are regions where the fROIs showed a reliable Hard > Easy Deduction effect in a left-out run; the ones indicated with black dots did not show this across-runs replicability.

**Supplementary Table 1.**
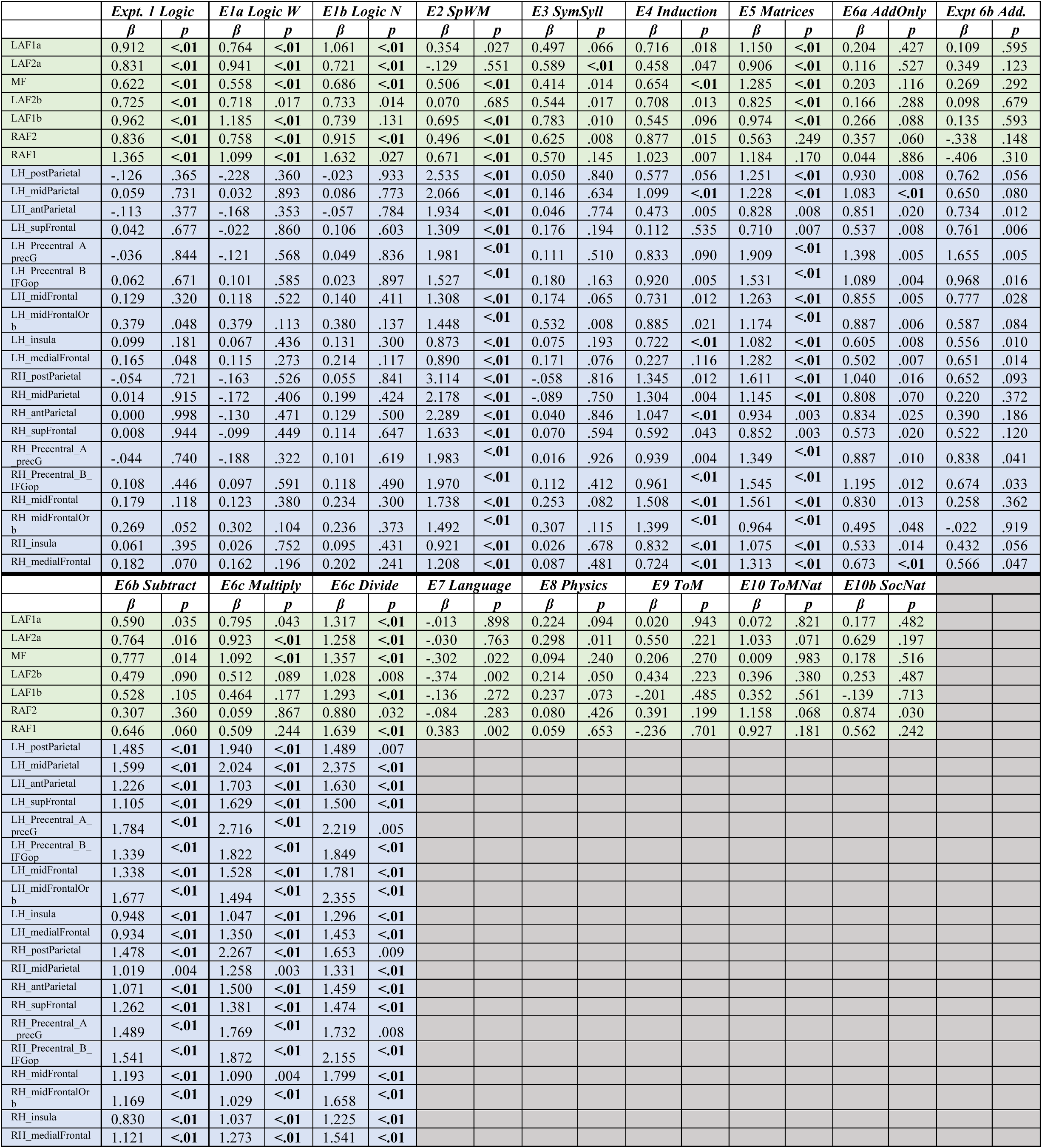
Individual Logic and MD fROIs’ responses to logical reasoning tasks and other tasks across Experiments 1-10.

**Supplementary Table 2.** Overlap between the logical-reasoning system and each of the other systems examined (the MD system, the language system, the physical-reasoning system and the ToM system), as measured with a Dice coefficient. We computed a Dice coefficient, (2 × |A ∩ B|) / (|A| + |B|), for each pair of contrasts for each participant at each of the four whole-brain thresholds. We report the averages across participants, and we also report a canonical non-overlap (between the Language and MD System) and the overlap between localizers for the MD System.

| Overlap between | Threshold: p < 0.0001 | Threshold: p < 0.001 | Threshold: p < 0.01 | Threshold: p < 0.05 |
| --- | --- | --- | --- | --- |
| MD1-MD1 | 0.402 | 0.452 | 0.497 | 0.525 |
| MD2-MD2 | 0.22 | 0.27 | 0.306 | 0.327 |
| MD1-MD2 | 0.287 | 0.321 | 0.364 | 0.406 |
| Logic-MD1 | 0.01 | 0.022 | 0.053 | 0.109 |
| Logic-MD2 | 0.012 | 0.025 | 0.057 | 0.103 |
| Logic-Phys | 0.007 | 0.022 | 0.06 | 0.117 |
| Logic-ToM | 0.048 | 0.053 | 0.073 | 0.113 |
| Logic-ToM2 | 0.014 | 0.03 | 0.061 | 0.102 |
| Logic-Soc | 0.002 | 0.008 | 0.032 | 0.075 |
| Logic-Lang | 0.006 | 0.012 | 0.03 | 0.066 |
| Lang-MD | 0.017 | 0.034 | 0.073 | 0.137 |

**Supplementary Table 3.** A. Comparison of Hard vs. Easy Deduction condition responses of the Logic Task in the MD and Logic Systems B. Comparison of Hard vs. Easy spatial WM condition responses of the Logic Task in the MD and Logic Systems.

| *A. Logical Syllogisms Task* |  |  |  |
| --- | --- | --- | --- |
| *Predictors* | *Beta* | *SEM* | *p* |
| Intercept [E Deduction] (in MD) | 2.32 | 0.28 | **<0.001** |
| H Deduction vs. E Deduction (in MD) | 0.07 | 0.07 | 0.347 |
| E Deduction (in MD) vs. E Deduction (Logic) | -1.47 | 0.38 | **<0.001** |
| H Deduction vs. E Deduction (in MD vs. in Logic) | 0.82 | 0.14 | **<0.001** |
| *B. Spatial Working Memory Task* |  |  |  |
| *Predictors* | *Beta* | *SEM* | *p* |
| Intercept [Easy SpWM] (in MD) | 1.84 | 0.33 | **<0.001** |
| H SpWM vs. E SpWM (in MD) | 1.49 | 0.08 | **<0.001** |
| E SpWM (in MD) vs. E SpWM (Logic) | -1.97 | 0.55 | **<0.001** |
| H vs. E (in MD vs. in Logic) | -1.11 | 0.16 | **<0.001** |

## Supplementary Methods

### Language localizer

Participants silently read sentences and lists of nonwords in a blocked design. The sentences > nonwords contrast targets cognitive processes related to high-level language comprehension, including understanding word meanings and combinatorial linguistic processing, and has been shown to effectively isolate language areas from nearby functional areas (see Fedorenko et al., 2024 for a review). The task is available for download from https://www.evlab.mit.edu/resources.

### MD network localizer (a spatial working memory task)

Participants are presented with 3x4 grids within which sequences of locations flash up (one at a time for a total of four locations in the easy condition, and two at a time for a total of eight locations in the hard condition). Participants are asked to keep track of the locations, and at the end of each trial, they are shown two sets of locations and asked to choose the set they just saw. The hard > easy contrast targets cognitive processes broadly related to performing demanding tasks—what is often referred to by an umbrella term ‘executive function processes’. The task is available for download from https://www.evlab.mit.edu/resources.

### Intuitive physical reasoning localizer

Participants are presented with videos of unstable block towers consisting of blue and yellow blocks (they are asked to judge which direction the tower would be most likely to fall in the intuitive physics condition, and whether the tower consists in more blue or yellow blocks in the color judgment condition). The physics > color judgment contrast targets cognitive processes related to intuitive physical reasoning. The task materials are available for download from https://osf.io/hqa23.

### Theory-of-Mind localizer

Participants read short vignettes about false beliefs (describing a false belief about the world in the mental representation condition, and otherwise matched vignettes about false or outdated world states in the physical representation condition) and answered a true-or-false comprehension question about the vignette. The mental > physical contrast targets cognitive processes related to reasoning about others’ or one’s own mental states—what is often referred to as “Theory-of-Mind” or social reasoning. The task is available for download from https://saxelab.mit.edu/use-our-efficient-false-belief-localizer.

### fMRI data acquisition

Whole-brain structural and functional data were collected on a whole-body 3 Tesla Siemens Trio scanner with a 32-channel head coil at the Athinoula A. Martinos Imaging Center at the McGovern Institute for Brain Research at MIT. T1-weighted structural images were collected in 176 axial slices with 1 mm isotropic voxels (repetition time (TR) = 2,530 ms; echo time (TE) = 3.48 ms). Functional, BOLD data were acquired using an EPI sequence with a 90° flip angle and using GRAPPA with an acceleration factor of 2; the following parameters were used: 31 4.4 mm thick near axial slices acquired in an interleaved order (with 10% distance factor), with an in-plane resolution of 2.1 × 2.1 mm, FoV in the phase encoding (A >> P) direction 200 mm and matrix size 96 × 96 voxels, TR = 2,000 ms and TE = 30 ms. The first 10 s of each run were excluded to allow for steady state magnetization.

### fMRI data preprocessing

fMRI data were preprocessed and analyzed using SPM12 (release 7487), CONN EvLab module (release 19b), and other custom MATLAB scripts. Each participant’s functional and structural data were converted from DICOM to NIFTI format. All functional scans were coregistered and resampled using B-spline interpolation to the first scan of the first session (Friston et al., 1995). Potential outlier scans were identified from the resulting subject-motion estimates as well as from BOLD signal indicators using default thresholds in CONN preprocessing pipeline (5 standard deviations above the mean in global BOLD signal change, or framewise displacement values above 0.9 mm; Nieto-Castañón, 2020). Functional and structural data were independently normalized into a common space (the Montreal Neurological Institute [MNI] template; IXI549Space) using SPM12 unified segmentation and normalization procedure (Ashburner and Friston 2005) with a reference functional image computed as the mean functional data after realignment across all timepoints omitting outlier scans. The output data were resampled to a common bounding box between MNI coordinates (−90, −126, −72) and (90, 90, 108), using 2 mm isotropic voxels and 4th order spline interpolation for the functional data, and 1 mm isotropic voxels and trilinear interpolation for the structural data. Last, the functional data were smoothed spatially using spatial convolution with a 4 mm FWHM Gaussian kernel.

### fMRI data modeling

For all experiments, effects were estimated using a general linear model (GLM) in which each experimental condition was modeled with a boxcar function convolved with the canonical hemodynamic response function (HRF) (fixation was modeled implicitly, such that all timepoints that did not correspond to one of the conditions were assumed to correspond to a fixation period). Temporal autocorrelations in the BOLD signal timeseries were accounted for by a combination of high-pass filtering with a 128 s cutoff, and whitening using an AR (0.2) model (first-order autoregressive model linearized around the coefficient a = 0.2) to approximate the observed covariance of the functional data in the context of restricted maximum likelihood estimation. In addition to experimental condition effects, the GLM design included first-order temporal derivatives for each condition (included to model variability in the HRF delays), as well as nuisance regressors to control for the effect of slow linear drifts, subject-motion parameters, and potential outlier scans on the BOLD signal.

### fROI definition and response estimation

#### Definition of the MD, language, physical, and social reasoning fROIs

Each individual map for the Critical > Control contrast from the relevant localizer was intersected with a set of parcels (5 left hemisphere language parcels, 20 MD parcels (10 per hemisphere), 19 Physics parcels (13 LH, 6 RH), and 10 ToM parcels) (5 per hemisphere). These parcels (available at OSF: https://osf.io/hqa23/) were derived from a probabilistic activation overlap map for the same contrast in a large set of independent participants (Language, n=220; MD, n=197; Physics, n=40; ToM, n=400). Within each parcel, a participant-specific fROI was defined as the top 10% of voxels with the highest t-values for the localizer contrast. To estimate the response in the fROIs to the conditions of the paradigm used to define the fROIs, the same cross-validation procedure was used as described above. As expected, all fROIs showed a robust Critical > Control effect (*p*s < 0.001, |d|s > 1.84; here and elsewhere, p-values are corrected for the number of fROIs using the false discovery rate (FDR) correction (Benjamini & Yekutieli, 2001)).

#### Functional correlation (a.k.a. functional connectivity) analysis

For each participant, the BOLD time-series was averaged across voxels within each fROI, and Pearson correlations were computed for every fROI pair (within the logic system, within the MD system, and across the two systems). Correlations were Fisher-transformed for normality, and differences among within- and between-system correlations were assessed with one-way ANOVAs. To visualize the results, Fisher z-values were averaged across participants to yield group-level fROI-to-fROI matrices (**Fig. 1E**), although all statistical analyses were performed within participants. We also performed a split-half reliability analysis by even- and odd-indexing our subjects and looking at the degree of similarity between the average BOLD response of the Logic and MD Systems of the even- vs. odd-indexed sets of subjects to the thirty-two syllogisms in the verbal deductive reasoning paradigm to check for the reliability of the response across subjects (**Fig. 1F)**.

#### Spatial correlations

For each participant, we computed a voxel-wise correlation between their whole-brain activation map for the Hard > Easy Deduction contrast and a map for each of the contrasts available for that participant. In addition, we computed a correlation within each contrast across runs. To equalize the amount of data for the between- vs. within-contrast comparisons, we also divided in half the data for the between comparisons, comparing odd-numbered runs for contrast 1 to even-numbered runs for contrast 2, then the opposite, and finally, averaging across the two to obtain a single value per participant per contrast pair. For visualization, Fischer-transformed correlations were averaged across participants.

#### Dice overlap analyses

For each participant, we computed a Dice overlap measure (Dice, 1945; Sørensen, 1948) between the thresholded activation maps for the Hard > Easy Deduction contrast and the Hard > Easy Spatial WM contrast (and other contrasts, for comparison). We used three whole-brain thresholds: p<0.001, p<0.01, and p<0.05 uncorrected; we included quite liberal thresholds to increase our chances of observing overlap if it exists.

**Supplementary Table 4.** Participant inclusion across experiments. Each row corresponds to an individual participant (UID), and each column indicates whether the participant contributed data to a given experiment. “Yes” marks inclusion in that experiment, and the bottom row lists the total number of participants per experiment. The experiments comprised Logic (Expt. 1), Spatial Working Memory (SpWM, Expt. 2), Symbolic Syllogisms (SymSyll, Expt. 3), Program Induction (ProgInd, Expt. 4), Cattell fluid matrix reasoning (Expt. 5), Math Addition only (Math_Add, Expt. 6a), Math with all four arithmetic operators (Math_Arith, Expt. 6b,c), Language (Lang, Expt. 7), Physics (Phys, Expt. 8), Theory of Mind (ToM, Expt. 9), and the naturalistic “Partly Cloudy” movie (Expt. 10).

|  | Logic | SpWM | SymSyll | ProgInd | Cattell | Math Add | Math Arith | Lang | Phys | ToM | ToM-nat |
| --- | --- | --- | --- | --- | --- | --- | --- | --- | --- | --- | --- |
| UID | Expt1 | Expt2 | Expt3 | Expt4 | Expt5 | Expt6a | Expt6b-c | Expt7 | Expt8 | Expt9 | Expt10 |
| 1065 | yes | yes | yes | yes | yes | yes | yes | yes | yes | yes | no |
| 1073 | yes | yes | yes | yes | yes | yes | yes | yes | yes | yes | no |
| 820 | yes | yes | yes | yes | yes | yes | yes | yes | yes | yes | yes |
| 1150 | yes | yes | yes | yes | yes | no | yes | yes | yes | yes | yes |
| 1151 | yes | yes | yes | yes | yes | no | yes | yes | yes | yes | yes |
| 1154 | yes | yes | yes | yes | yes | no | yes | yes | yes | yes | yes |
| 1157 | yes | yes | yes | yes | yes | no | yes | yes | yes | no | yes |
| 1177 | yes | yes | yes | no | yes | no | yes | yes | yes | yes | yes |
| 1229 | yes | yes | yes | no | yes | yes | no | yes | no | no | yes |
| 1248 | yes | yes | yes | no | yes | no | yes | yes | no | no | no |
| 1257 | yes | yes | yes | no | yes | no | no | yes | no | no | yes |
| 1268 | yes | yes | yes | no | yes | no | no | yes | no | no | no |
| 1250 | yes | yes | yes | no | yes | no | no | yes | no | no | no |
| 1249 | yes | yes | yes | no | no | no | no | yes | no | no | no |
| 1080 | yes | yes | no | yes | no | yes | yes | yes | yes | yes | yes |
| 1074 | yes | yes | no | yes | no | yes | no | yes | yes | yes | yes |
| 1069 | yes | yes | no | yes | no | yes | no | yes | yes | yes | no |
| 1016 | yes | yes | no | yes | no | yes | no | yes | yes | no | no |
| 1059 | yes | yes | no | yes | no | yes | no | yes | yes | no | no |
| 1071 | yes | yes | no | yes | no | yes | no | yes | yes | no | no |
| 865 | yes | yes | no | yes | no | no | yes | yes | yes | no | yes |
| 892 | yes | yes | no | yes | no | no | yes | yes | yes | no | no |
| 1147 | yes | yes | no | yes | no | no | yes | yes | yes | no | no |
| 1152 | yes | yes | no | yes | no | no | yes | yes | yes | no | no |
|  | *n=24* | *n=24* | *n=14* | *n=17* | *n=13* | *n=10* | *n=14* | *n=24* | *n=18* | *n=10* | *n=11* |

## Notes

https://osf.io/hqa23

### Competing Interest Statement

The authors have declared no competing interest.

